# Does hybridisation with an invasive species threaten Europe’s most endangered reptile? Genomic assessment of Aeolian lizards on Vulcano island

**DOI:** 10.1101/2024.04.12.589112

**Authors:** Josephine R Paris, Gentile Francesco Ficetola, Joan Ferrer Obiol, Iolanda Silva- Rocha, Miguel Angel Carretero, Daniele Salvi

**Affiliations:** Department of Health, Life and Environmental Sciences, University of L’Aquila, Coppito, Italy; Department of Life and Environmental Sciences, Marche Polytechnic University, Ancona, Italy; Department of Environmental Science and Policy, University of Milan, Milano, Italy; Université Grenoble Alpes, Université Savoie Mont Blanc, CNRS, Laboratoire d’Ecologie Alpine (LECA), Grenoble, France; CIBIO, Centro de Investigação em Biodiversidade e Recursos Genéticos, Universidade do Porto, InBio Laboratorio Associado, Vairão, Portugal; Departamento de Biologia, Faculdade de Ciências da Universidade do Porto, Porto, Portugal; BIOPOLIS Program in Genomics, Biodiversity and Land Planning, CIBIO, Campus de Vairão, 4485-661 Vairão, Portugal

**Keywords:** endangered, endemic, hybridisation, islands, lizards, *Podarcis raffonei*, *Podarcis siculus*, RADseq

## Abstract

Interspecific hybridisation can be consequential for rare and insular endemic species. The Critically Endangered Aeolian wall lizard, *Podarcis raffonei*, severely declined due to interactions with the invasive Italian wall lizard, *Podarcis siculus*. The largest population of *P. raffonei* survives on a narrow peninsula (Capo Grosso) that is mildly connected to the island of Vulcano, which has been entirely invaded by *P. siculus*. Recent observation of individuals with an intermediate phenotype raised concern over the risk that hybridisation might swamp this last stronghold. We genetically characterised lizards from Vulcano using genome-wide SNPs, considering individuals showing multiple phenotypes (native, invasive, and “intermediate”). Hybridisation rate was low (∼3%), with just two F1 hybrids and two backcrosses. However, pure *P. raffonei* showed extremely low genetic diversity, a very small effective population size, and a low N_E_/N_C_ ratio. Management strategies are urgently needed to control invasive species and maintain the genetic diversity of *P. raffonei*.

## 1. Introduction

Climate change, habitat modification, and the intentional or unintentional movement of organisms by humans, are causing species distribution shifts across the planet. One outcome of these disturbances is increased rates of interspecific hybridisation, which can be especially problematic for rare or endangered species that encounter more abundant species (Rhymer & Simberloff, 1996; Allendorf *et al*., 2001). Hybridisation can threaten a species’ genetic integrity and even contribute to species extinction (Levin *et al.,* 1996; Rhymer & Simberloff, 1996; Allendorf *et al*., 2001), yet in some cases hybridisation can also increase diversity and may aid evolutionary rescue of endangered species (Stelkens *et al*., 2014; Hoffmann *et al*., 2015; Vedder *et al*., 2022). Nonetheless, the effects of interspecific hybridisation are highly species– and context-specific, and to predict its potential evolutionary outcomes, a robust quantification of hybridisation is required. Such evaluations are especially pressing for rare endangered species, where an understanding of the rate and mode of interspecific hybridisation is required to design the most appropriate conservation and management plans.

Alien invasive species are a major cause of extinction risk in reptiles, with particularly strong effects in island environments (Böhm *et al*., 2013; Spatz *et al.,* 2017; Cox *et al.,* 2022). In this respect, the endemic Aeolian wall lizard, *Podarcis raffonei*, provides an emblematic case. Once probably widespread across the Aeolian archipelago (Southern Italy), this lizard is now restricted to three tiny islets and a narrow peninsula (Figure 1). With an estimated range <0.02 km^2^, *P. raffonei* is the rarest and most threatened European reptile. Listed as Critically Endangered by the IUCN since 2006, the species remains understudied and underprotected (Gippoliti *et al*., 2017). The island of Vulcano hosts the only population of *P. raffonei* that is not microinsular as it occurs on a narrow peninsula (Capo Grosso, surface < 0.7 ha) connected to a main island. Until 1999, the species was recorded in multiple locations across Vulcano island (Vulcanello and Gran Cratere) (Capula *et al*., 2002), but these populations are now considered to be extinct (Lo Cascio, 2010; D’Amico *et al*., 2018; Lo Cascio & Sciberras, 2020). Interspecific interactions with the Italian wall lizard, *P. siculus*, have been regarded as a major cause of the near-extinction of *P. raffonei* (Capula *et al*. 2002). The Italian wall lizard is native of the Italian Peninsula, Sicily, and the northern Adriatic coast, but has been introduced by humans across the world (Silva-Rocha *et al*., 2014; Bonardi *et al*., 2022). *P. siculus* has been likely introduced in Vulcano and other Aeolian islands a few centuries ago (Sherpa *et al*., 2023) and is now widespread across the archipelago (Capula *et al*., 2002; Lo Cascio, 2010; D’Amico *et al*., 2018). The invasion by *P. siculus* has thus been proposed as a key driver for *P. raffonei’*’s relict distribution (Capula, 1993, 1994; Capula *et al*. 2002).

**Figure 1.**
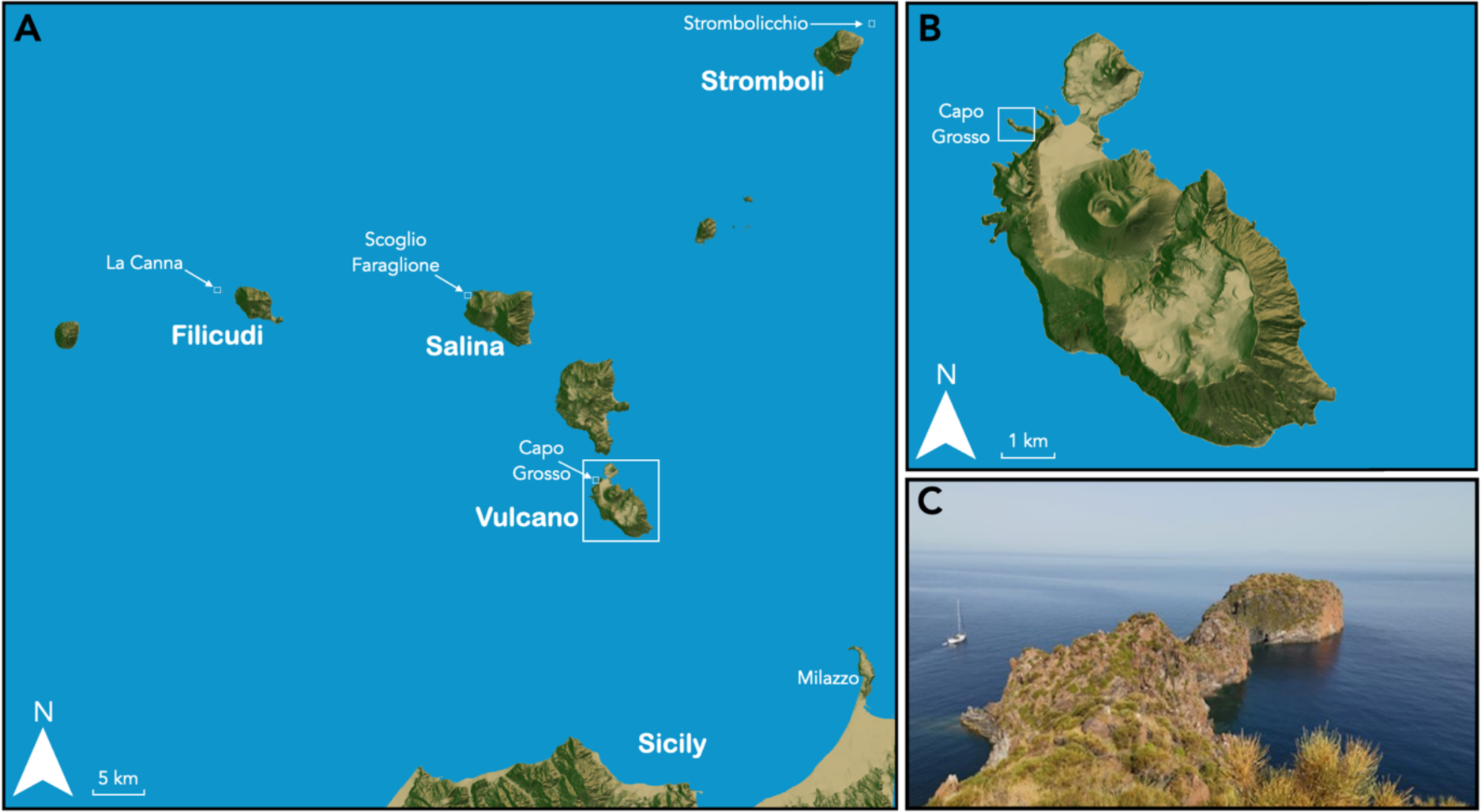
Geographic locations of the Aeolian wall lizard (*Podarcis raffonei*), sampling locations. (**A**) Map of the Aeolian Islands in southern Italy. The locations of the Aeolian wall lizard (*Podarcis raffonei*) are the three islets: La Canna (close to Filicudi), Scoglio Faraglione (close to Salina), Strombolicchio (close to Stromboli) and the peninsula of Capo Grosso on the island of Vulcano. Sampled individuals include: the typical brown phenotype (*P. raffonei;* n=50) and putative hybrid lizards with an intermediate phenotype (n=38) from Capo Grosso; pure *P. raffonei* individuals from Scoglio Faraglione (n=5); and pure *P. siculus* lizards from mainland Vulcano (n=35) and from mainland northern Sicily, Milazzo (n=5). (**B**) Zoom-in on the island of Vulcano, showing the small peninsula of Capo Grosso. (**C**) Photograph of Capo Grosso: *P. raffonei* is confined to the small distal portion of the peninsula. Orographic maps were created in R v4.0.2 using rayshader v0.28.2 (Morgan-Wall, 2022).

Italian wall lizards might impact Aeolian wall lizards through both competition and hybridisation. Competition has been proposed as a reason for decline because *P. siculus* has a larger body size, higher aggressiveness, excellent thermoregulation, behavioural plasticity, and tolerance to disturbed environments (Capula 1992; Capula *et al*., 2002; Downes & Bauwens 2002; Damas-Moreira *et al*., 2018, 2019; Caruso 2021). Moreover, an early allozyme analysed 101 lizards from Vulcano, identifying 15 of them as F1 hybrids between *P. raffonei* and *P. siculus* (hybrid ratio: ~ 15%; Capula, 1993), suggesting that hybridisation might further accelerate the decline of *P. raffonei* (Capula, 1993; Capula *et al*., 2002; Ficetola *et al*., 2021). However, these early studies only used a small panel of allozyme markers (only four “diagnostic” loci; Capula, 1993). The assessment of hybridisation patterns requires very large (>100) panels of markers, as a smaller number of markers can yield both severe underestimation and overestimation of hybridisation rates (Ravagni *et al.,* 2021). Unfortunately, such assessments are so far lacking.

More recently, observations of lizards from Capo Grosso with a green dorsal-colouration phenotype (hybrid-like) resulted in renewed alarm (Ficetola *et al*., 2021). *P*. *raffonei* and *P. siculus* differ only slightly in morphology and colour pattern, meaning the two species are difficult to tell apart (Capula, 1993; Capula & Lo Cascio 2011). *P. raffonei* possesses dark markings on the throat and is generally brown on the dorsal surface, whereas *P. siculus* has a white throat with no dark spots and shows a green dorsal pattern. In a survey of 131 lizards in 2017, a large number (>50%) of individuals from Capo Grosso showed an intermediate phenotype, with green patterns and dark markings on the throat (Figure 2). This was suspected to indicate extensive hybridisation (Ficetola *et al.,* 2021); however, individuals kept in captive-breeding programs showed strong plasticity in colour pattern with a seasonal shift towards a green phenotype (Ficetola *et al.,* 2021). Thus, the true hybrid status of the Capo Grosso population of *P. raffonei* remains unknown. Given the high-priority conservation status of the Aeolian wall lizard, and the potentially catastrophic effects of widespread hybridisation with the Italian wall lizard, there is a pressing need to quantify the actual hybridisation rate using a large panel of genomic markers. Such assessment would also provide the first measures of genetic diversity and effective population size (N_E_) of the largest extant population of *P. raffonei*, with key consequences for the management of this island endemic.

**Figure 2.**
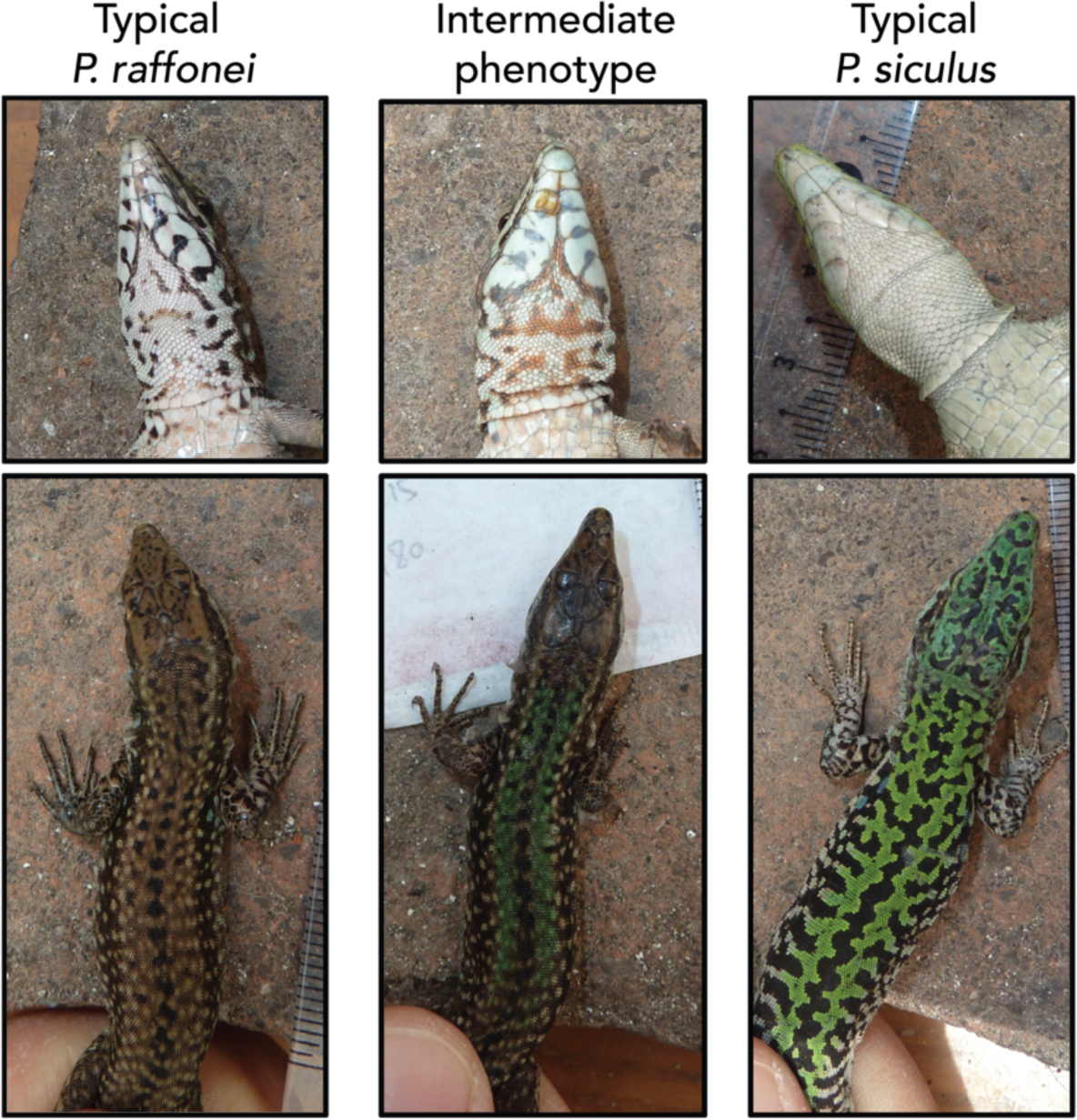
Typical colour patterns of males of the Aeolian wall lizard *Podarcis raffonei* (left) and the Italian wall lizard *P. siculus* (right), and patterns of individuals with the intermediate phenotype (middle) from Ficetola *et al*. (2021). Typical phenotypes of *P. raffonei* and intermediate phenotypes were observed on the Capo Grosso peninsula, whereas typical phenotypes of *P. siculus* were observed across the main island of Vulcano.

In this study, we use single-nucleotide polymorphisms (SNPs) derived from double-digest restriction-site associated DNA sequencing (ddRAD-seq) to investigate the status of *P. raffonei* from the Capo Grosso peninsula. First, we quantified the genetic status of lizards from Capo Grosso and mainland Vulcano, evaluating the extent of hybridisation between *P. raffonei* and *P. siculus*, and assessing the relationship between genotype and colour pattern. Second, after the exclusion of hybrids, we performed an assessment of genetic diversity and effective population size (N_E_) of *P. raffonei* from Capo Grosso. These results have key implications for the conservation and management of this Critically Endangered species, and provide general insights on the conservation of small, isolated populations under the increasing threat of hybridisation with invasive species.

## 2. Materials and Methods

### 2.1 Study species and population

*Podarcis raffonei* occurs in three isolated, tiny islets (Scoglio Faraglione, Strombolicchio and La Canna), and a narrow peninsula (Capo Grosso) on the island of Vulcano (Fig. 1). Today, Capo Grosso (surface < 0.7 ha) is the only known location of *P. raffonei* on Vulcano and represents more than half of the remaining range and of this species (Ficetola *et al*., 2021). Moreover, Capo Grosso represents the only remaining population of *P. raffonei* that is still connected to a main island.

### 2.2 Study sites and sampling

Sampling occurred during the spring and summer of 2015 and 2017 and comprised a total of 138 lizards (Table S1). Adult lizards were captured with a noose. For each lizard, we recorded the sex (based on sexual secondary characters and hemipenis eversion) and took standard photographs of dorsal and ventral patterns. A 2cm tail clip was obtained and stored in 95-100% EtOH for genetic analysis. In Capo Grosso, we sampled 54 lizards with the brown phenotype typical of *P. raffonei* and 38 lizards with a green dorsal-colouration phenotype (hereafter “intermediate”), matching the description of hybrids reported in previous studies; Capula *et al.,* 1993; Capula & Lo Cascio 2011). Furthermore, we sampled 36 lizards from the main island of Vulcano, matching the typical phenotype of *P. siculus* (Fig. 1 & 2). To ensure the correct identification of each species group from the Capo Grosso/Vulcano system, we also sampled five pure *P. raffonei* and five pure *P. siculus* from two locations where the two species do not overlap: *P. raffonei* from the islet of Scoglio Faraglione and *P. siculus* from Milazzo (mainland Sicily).

### 2.3 DNA extraction and ddRAD library preparation

Genomic DNA was extracted using the DNeasy Blood & Tissue kit (Qiagen, Germany), following manufacturer guidelines. ddRAD library preparation included an initial digestion of 300 ng of DNA in a 34 μL reaction (2 hours at 37 °C; 10 U each of *SbfI* and *MspI*, New England Biolabs Inc.). Standard Illumina adapters were ligated using 60 cycles of digestion at 37 °C (2 minutes) and ligation at 16 °C (4 minutes) with 400 U of T4 ligase (New England Biolabs, USA), followed by heat-inactivation at 65 °C (10 minutes). Digested-ligated products were purified using 1:5:1 ratio of AMPure XP beads (Beckman Coulter, USA). Size selection was performed using a BluePippin to retain total fragment sizes of 250-500bp and were purified using the QIAquick Gel Extraction Kit (Qiagen, Germany). Libraries were obtained by pooling 10 x 20 μL of the each PCR reaction per library, each consisting of 2.5 μL of DNA, 0.2 mM of dNTPs, 0.15 μM of primer, 3% dimethyl sulphoxide (DMSO) and 0.4 U of Taq-Phusion High-Fidelity (New England Biolabs, USA). PCR conditions were as follows: initial denaturation at 98 °C (10 minutes), 12 cycles of 98 °C (10 s), 66 °C (30 s), 72 °C (1 min), final extension period at 72 °C (10 min). Libraries were purified with QIAgen MinElute PCR Purification Kit (Qiagen, Germany) and were sequenced on an Illumina HiSeq 2500 (2 x 125 bp).

### 2.4 Data processing

Reads shorter than 125 bp in length were removed using *fastp* v0.23.2 (Chen *et al*., 2018). Stacks v2.60 (Rochette *et al*., 2019) was used to assemble loci. Trimmed data were cleaned and demultiplexed using the module *process_radtags*. For hybrid detection, we used a *de novo* approach to avoid biassed estimations of allele frequencies by aligning to a single species’ genome. *P. raffonei* and *P. siculus* were first assembled separately to remove low quality samples (Cerca *et al*., 2021) and to perform parameter optimisation (Paris *et al.,* 2017). Due to the high divergence time between the species (11-18 million years; Salvi *et al*., 2021; Yang *et al*., 2021), parameters were optimised for each species separately in *ustacks* (M parameter). We then optimised to species-specific loci into a catalog by assessing the n parameter across species. The catalog for *de novo* optimisation consisted of loci assembled from 20 individuals with the highest coverage from each of pure *P. raffonei* and pure *P. siculus.* For estimates of genetic diversity and effective population size, we used RAD loci aligned to the reference genome of *P. raffonei* (GCF_027172205.1; Gabrielli *et al*., 2023). Cleaned reads were aligned to the genome using bwa mem v0.7.17 (Li, 2013), marking shorter split hits as secondary reads (–M option). Secondary reads were removed from the alignment files prior to processing with *ref_map*. As different analyses require differently filtered datasets, we created several subsets from our data using the *populations* module. Further details of these filters can be found below in each relevant section.

### 2.5 Genetic structure and hybrid identification

The *de novo* assembled loci were used for analyses of genetic structure and hybrid identification. We created a whitelist of loci present only in *P. siculus* green and *P. raffonei* brown individuals (no intermediate phenotypes), including the pure known individuals from Scoglio Faraglione (*P. raffonei*) and Milazzo (*P. siculus*). Loci had to be present in both groups (–p 2), and in 50% of individuals from each of these groups (–r 0.5), including alleles present at a minor allele count of 2 (––min-mac 2). This whitelist was used to sample loci across all individuals, inclusive of the intermediate-phenotype samples. The dataset was then assessed for depth in vcftools v0.1.17 (Danecek *et al*., 2011) and was filtered accordingly (––minDP 3 and ––max-meanDP 80), keeping only biallelic sites at a genotype quality of 30, resulting in a linkage-pruned dataset (––write-single-SNP) comprised of 2,623 variant sites. The same procedure was used to generate data for the full haplotype (all SNPs for each RAD locus), resulting in a dataset comprising 17,254 variant sites. Data manipulation and plotting was performed in R v4.0.2 with ggplot2 (Wickham, 2016) and tidyverse (Wickham *et al.,* 2019).

A Principal Components Analysis (PCA) was performed using the linkage-pruned dataset in Plink v1.9 (Purcell *et al*., 2007). Pairwise-*F*_ST_ between the two species was calculated using the linkage-pruned dataset in Hierfstat v0.5.11 (Goudet, 2005). The full haplotype information was used for high-resolution inference of the recent shared ancestry between the samples using fineRADstructure v0.3.2 (Mendes *et al*., 2018), running the model for 100,000 MCMC iterations with a thinning interval of 1000, discarding the first 10,000 iterations as burn-in, and building a tree with 10,000 hill-climbing iterations. We categorised fixed loci from individuals showing the highest differences in PCA space (36 *P. siculus* samples and 55 brown *P. raffonei* individuals), resulting in 1081 markers. Note that samples with an intermediate phenotype were not used for marker selection. These fixed markers were used to estimate hybrid indices in introgress v1.2.3 (Gompert & Buerkle, 2010). Hybrid indices and their confidence limits were estimated using the *est.h* function using 1000 bootstraps.

Admixture v1.3 (Alexander *et al.,* 2009) was used for maximum-likelihood (ML) estimates of individual ancestries using the linkage-pruned dataset. To assess variability in ML estimates, the algorithm was run ten times, with the number of groups (*K*) varying between 1 and 7 with a tenfold cross-validation (CV=10). After identification of the optimal *K*, we performed a hierarchical analysis to identify further sub-structuring within the groups using Admixture. We also performed admixture analysis to assess whether there were any biases introduced into the data by sampling year (2015 versus 2017).

The posterior probability of each intermediate-phenotype individual belonging to one of each discrete genotype classes (pure *P. siculus*, pure *P. raffonei*, F1 hybrid, F2 hybrid, *siculus* backcross and *raffonei* backcross) was estimated using NewHybrids v2.0 (Anderson & Thompson, 2002). Data were analysed in parallel using *parallelenewhybrid* (Wringe *et al*., 2017a) as implemented in *hybriddetective* (Wringe *et al*., 2017b). All analyses were performed with 200,000 MCMC sweeps, discarding the first 50,000 sweeps as burn-in, using Jeffries-like priors for both the allele frequency (*θ*) and mixing proportion (π). Genotype frequency classes were simulated using a panel of loci derived from brown phenotype *P. raffonei* from Capo Grosso, *P. raffonei* from Scoglio Faraglione, *P. siculus* from Vulcano and *P. siculus* from Milazzo. Based on our empirical data, we simulated n=50 of each pure class (*P. siculus* or *P. raffonei*), n=5 F1s, n=3 F2s, n=2 *siculus* backcross and n=2 *raffonei* backcross using the HybridLab algorithm (Nielsen *et al*., 2006). We simulated three independent datasets, each of which was analysed three times. To evaluate marker efficiency, we explored three random subsets of the 50 most-informative markers (high *F*_ST_, linkage-pruned). Convergence of the simulations was checked by assessment of the critical posterior probability of assignment thresholds to each genotype frequency class. Pure individuals from the simulated data were then combined with empirical data for the intermediate phenotype individuals. The z flag was assigned to the pure genotype frequency categories.

Finally, we created a whitelist containing RAD loci genotyped across all individuals (*i.e.*, no missing data) which were alternatively fixed between pure *P. siculus* and pure *P. raffonei* (*F*_ST_ =1) to explore the signature of individual genotypes. This analysis was conducted in attempts to better clarify the hybrid status of the low coverage individual (CGI38). The stringent filtering resulted in 87 SNPs genotyped across 49 marker loci. Genotypes were plotted using the Genotype Plot function in R (Whiting, 2022).

### 2.6 Characterising the parental sex and phenotypes of the genetic hybrids

As mitochondria are inherited in a matrilinear way, a hybrid lizard contains the mitochondrial DNA (mtDNA) of its ‘mother species’. The maternal species of each of the identified hybrid individuals was therefore inferred by sequencing a fragment of the 12S rRNA mtDNA region using PCR primers and protocols described in Mendes *et al.,* (2016). We characterised the overall phenotype of hybrids by inspecting the distribution of characteristics typical of the two parental species as reported in literature (see Capula & Lo Cascio 2011) for *P. raffoeni*: colour of back generally brown (greener in spring) with small dark dots often aligned in a vertebral stripe, the presence of evident dark markings on the throat (usually absent in *P. siculus*); and for *P. siculus*: extensive green in the back with a striped or reticulated pattern, a uniform white throat and belly.

### 2.7 Estimating the genetic diversity and effective population size of *P. raffonei* in Capo Grosso

For estimates of genetic diversity, we included only individuals sampled in 2017. Loci were filtered so that they had to be present in both *P. raffonei* and *P. siculus* (–p 2) and present in all individuals (–r 1) at a minor-allele count of 2 (––min-mac 2), and loci mapping to the Z and W chromosome were excluded (total loci = 26,831). To account for any potential artefacts due to missing data, we estimated genetic diversity based only on individuals with low amounts of missing data (*P. raffonei* = n60; *P. siculus* = n30). As the *P. siculus* dataset comprised half the number of individuals, we evaluated any potential sample size bias by randomly downsampling the *P. raffonei* dataset to the same number of individuals as the *P. siculus* dataset (n=30). Observed heterozygosity (H_0_), expected heterozygosity (H_E_), the inbreeding coefficient (F_IS_) and allelic richness (A_R_) were calculated in Hierfstat v 0.5.11 (Goudet, 2005). The *F*_IS_ values were bootstrapped over loci 1000 times to obtain 95% confidence intervals. Allelic richness estimates were rarefied against a sample size of 20 diploids. Estimates for nucleotide diversity (π) were derived from the Stacks *populations* output.

Estimates of the effective population size (N_E_) were calculated using the linkage disequilibrium (LD) method implemented in NeEstimator v2.1 (Do *et al*., 2014). Estimates were only calculated for the identified true *P. raffonei* lizards from Capo Grosso (n=74) with no missing data (6,772 loci). The *P. siculus* dataset comprised too few individuals to infer accurate estimates. We excluded the Z and W chromosomes from analysis. The N_E_ was estimated both with and without singletons and 95% confidence limits were calculated using jack-knifing, using a *Pcrit* value = 0.05. Resulting effective population size estimates were used to calculate the N_E_/N_C_ ratio by using *N*-mixture model census size (N_C_) estimates for the Capo Grosso population: 1050 (847 – 1280) (Ficetola *et al*., 2021).

## 3. Results

### 3.1 ddRAD data quality control, *de novo* assembly and reference alignment

After removal of five low-quality samples, we generated over 224 million paired-end reads (average 1,835,851 ± SE 85,979 per individual) for 133 lizards: Capo Grosso brown phenotype (n=50); Capo Grosso intermediate phenotype (n=38); Vulcano *P. siculus* (n=35); Scoglio Faraglione *P. raffonei* (n=5); Milazzo *P. siculus* (n=5) (Table S1). *De novo* assembly of the RAD data separately for *P. raffonei* and *P. siculus* showed that the best parameter for merging alleles into loci (–M) was 2 for both species. We merged the loci across individuals from both species (–n) using an optimised value of 2. Average coverage overall was 31X. One individual (CGI38) was retained for analysis, despite low coverage (5.4X) as it showed evidence of being a hybrid in a preliminary assessment of the dataset. In the reference-guided analysis, *P. raffonei* and *P. siculus* samples showed different alignment rates to the *P. raffonei* reference genome, with *P. raffonei* samples showing an average alignment rate of 92.5%, and *P. siculus* samples showing an average alignment rate of 72% (Table S1). This alignment bias was alleviated by retaining loci occurring in both species (–p 1 in *populations*).

### 3.2 Inter– and intra-species genetic structure and identification of genetic hybrids

PCA showed strong genetic differentiation between *P. siculus* lizards sampled from Vulcano and brown *P. raffonei* individuals sampled from Capo Grosso (Figure 3A). PC1 (87% of the variance) clearly separated the two species and evidenced that 34 of the intermediate phenotype lizards sampled from Capo Grosso clustered with the brown *P. raffonei* lizards. Four of the intermediate-phenotype lizards were placed in the genetic space between the two species: CGI13 and CGI14 appeared in the middle of PC1; CGI38 clustered approximately in the middle, but closer to the *P. siculus* cluster; and CGI36 was closer to the *P. raffonei* cluster. PC2 (1% of the variance) separated the *P. raffonei* lizards sampled from Capo Grosso from lizards sampled from Scoglio Faraglione. *P. siculus* lizards sampled from Milazzo showed very little genetic differentiation from *P. siculus* from Vulcano, as indicated by PC3 (0.93% of the variance). Genetic differences were well-reflected in the *F*_ST_ estimates among the four sampling locations (excluding intermediate phenotypes). High *F*_ST_ was observed between brown *P. raffonei* lizards from Capo Grosso and *P. siculus* from Vulcano (*F*_ST_ = 0.92). High *F*_ST_ was also found between *P. raffonei* lizards from Capo Grosso and *P. raffonei* from Scoglio Faraglione (*F*_ST_ = 0.42). Low *F*_ST_ was observed between *P. siculus* lizards from Vulcano and Milazzo (*F*_ST_ = 0.07).

**Figure 3.**
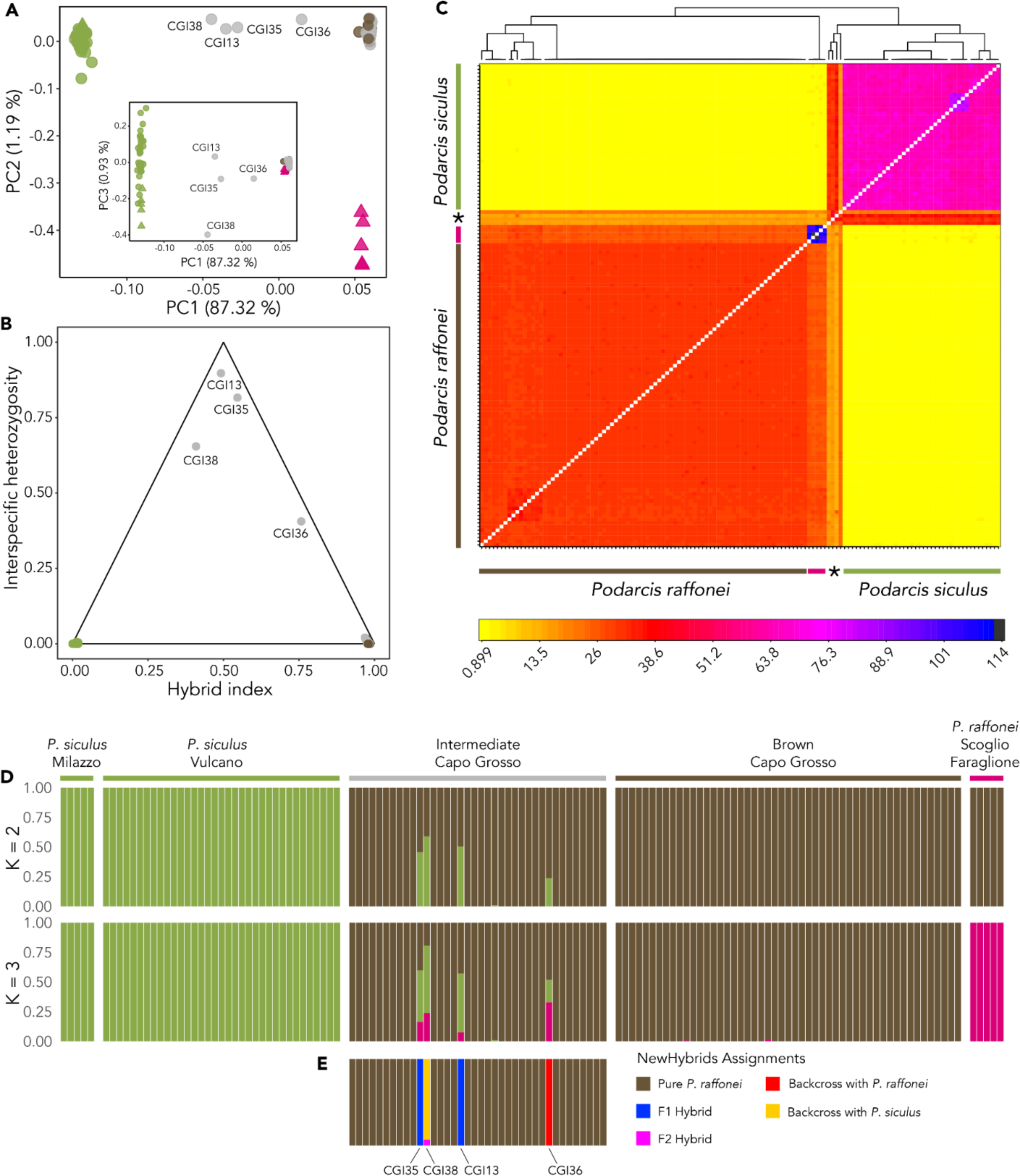
Genetic structure and identification of genetic hybrids. A) Principal Components Analysis (PCA Green circles represent *P. siculus* lizards from Vulcano; green triangles represent *P. siculus* lizards from Milazzo; brown circles represent brown-phenotype lizards from Capo Grosso; pink triangles represent *P. raffonei* lizards from Scoglio Faraglione; grey circles represent intermediate-phenotype lizards from Capo Grosso. Main plot shows the variance explained by PC1 and PC2; inset shows the variance explained by PC1 and PC3. B) Triangle plot depicting the hybrid index, and heterozygosity at species diagnostic SNPs inferred by introgress. Green represents *P. siculus* lizards; brown represents brown-phenotype lizards, grey represents intermediate-phenotype lizards. C) Co-ancestry relationships from fineRADstructure. Two main clusters are observed, representing *P. siculus* and *P. raffonei*. All intermediate-phenotype lizards (except the four hybrids, identified by the asterisk) cluster within the *P. raffonei* cluster. *P. siculus* from Milazzo are within the *P. siculus* cluster. *P. raffonei* lizards from Scoglio Faraglione show a distinct cluster within the *P. raffonei* cluster of Capo Grosso. Colour legend depicts the proportion of shared ancestry between the haplotypes. The tree atop shows crude relationships between the haplotypes. D) Individual ancestries from Admixture showing results for *K*=2 and *K*=3. E) Assignment of the intermediate-phenotype lizards using NewHybrids. The colour legend describes the assignment to each of the five genotype classes that were identified.

Introgress analysis confirmed that most intermediate-phenotype lizards are in fact *P. raffonei*, with the analysis indicating just four genetic hybrids based on the hybrid index and interspecific heterozygosity (Figure 3B). CGI13 (male) and CGI35 (female) showed F1 hybrid indices of approximately 50% [CGI13: 0.492 (0.467 – 0.512); CGI35: 0.546 (0.523 – 0569)]. CGI36 (male) had a back-cross hybrid index: 0.758 (0.737 – 0.778). The low-coverage individual (CGI38; male) showed a hybrid index more closely related to *P. siculus*, although with larger 95% confidence intervals: 0.409 (0.344 – 0.476).

The co-ancestry method of fineRADstructure using the full haplotype information also showed two clearly defined clusters representing the two species (Figure 3C). *Podarcis raffonei* from Scoglio Faraglione clustered within the *P. raffonei* cluster, and *P. siculus* from Milazzo clustered within the *P. siculus* cluster, each with higher co-ancestries to the other samples from these sampling locations. The fineRADstructure analysis further confirmed the identification of the four hybrids. The two suspected F1 hybrids (CGI13 and CGI35) showed equally shared donor and recipient ancestries to both *P. siculus* and *P. raffonei*. Sample CGI36 showed higher shared co-ancestry to *P. raffonei*. Sample CGI38 (low coverage) showed approximately equally shared donor and recipient ancestries to both species, although with some small haplotype blocks showing a stronger signature of donor ancestry from *P. siculus*.

Cross-validation (CV) error from Admixture showed that *K* = 3 was the best *K* for describing the allocation of individual ancestries, but *K* = 2 also showed a low CV error rate (Figure S1). At *K* = 2, individuals from the two species clustered into two clearly defined groups (>99% assignment probability (Figure 3D). Most of the intermediate phenotype samples showed the same cluster assignment (>99%) as the brown-phenotype *P. raffonei* from Capo Grosso and *P. raffonei* from Scoglio Faraglione. Four putative hybrids were clearly apparent from their shared assignment probabilities to the two species clusters. Two samples showed approximately 50% assignment probability to *P. raffonei* (CGI13: 50%; CGI35: 54%). Sample CGI36 showed a high assignment probability to *P. raffonei* (CGI36: 0.76%). CGI38 (low coverage) showed a higher assignment to *P. siculus* (CGI38; 59%). At *K* = 3, the *P. raffonei* samples from Scoglio Faraglione were separated from the Capo Grosso *P. raffonei* cluster and the four hybrid individuals also showed some shared ancestry with this location. Further assessment of population structure within the groups showed that the best *K* was 1 for both the Capo Grosso *P. raffonei* individuals and the Vulcano *P. siculus* individuals, and there was no effect of sampling year (Figure S2). Hierarchical analysis on all *P. siculus* samples also showed *K* = 1 as the best value for *K*, although examination of the cluster probabilities for *K* = 2 showed that lizards sampled from Milazzo formed a homogenous group compared to samples from Vulcano (Figure S3).

In the NewHybrids assignment, all panels of diagnostic markers used for simulation were species-diagnostic (*i.e.*, *F*_ST_ = 1). The critical posterior probability of assignment thresholds to each genotype frequency class was 1 for all marker panels across all runs in the simulated datasets. We combined the empirical data with all three marker panels to assess hybrid classification of the intermediate-phenotype individuals. These results also confirmed that the majority (n=34) of individuals with the intermediate phenotype are *P. raffonei* (Figure 3E). NewHybrids assigned two of the intermediate-phenotype samples as F1 hybrids (CGI13 and CGI35), and one sample as a *P. raffonei* backcross (CGI36) with high posterior probabilities (>0.99 across all runs). The low-coverage sample (CGI38) showed the highest assignment probability of being a *siculus* backcross. However, the reduced data quality of this sample resulted in a lower assignment probability across all marker panels and simulations (0.941; range: 0.940 – 0.942), with the remaining assignment probability assigning this individual as a F2 hybrid.

Finally, we used a whitelist of 87 SNPs genotyped across 49 loci to interrogate the individual genotype information of the four identified hybrid individuals (Figure 4). This confirmed the genotype patterns for the four genetic hybrids, with CGI13 and CGI35 showing heterozygous *P. raffonei* and *P. siculus* genotypes for the SNPs used in the analysis. CGI36 had a majority of homozygous *P. raffonei* genotypes, but a stretch of heterozygous genotypes was also seen, as would be expected with a *raffonei* backcross. In the low-coverage individual (CGI38), a large proportion of the SNPs were heterozygous, but a genotype signature of homozygous *P. siculus* was also observed.

**Figure 4.**
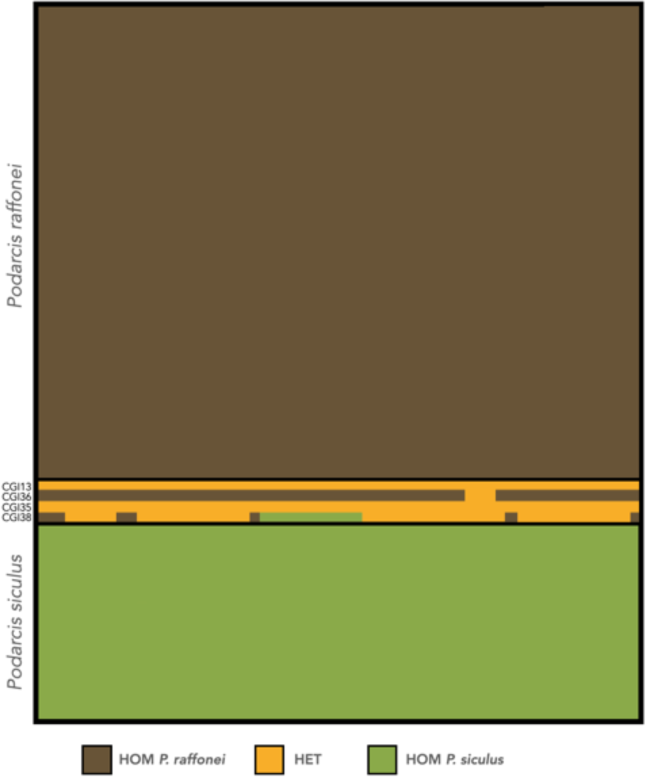
Genotypes plots for 87 species-diagnostic SNP markers (*i.e. F*_ST_ = 1) between *P. raffonei* and *P. siculus*. Genotypes are coloured as being homozygous in *P. raffonei* (HOM = brown), homozygous in *P. siculus* (HOM = green) or heterozygous between the two species (HET = yellow). Each block is coloured by these genotype calls. Note that the SNPs were called *de novo* and are therefore not ordered in a positional manner.

This further confirms that the genetic hybrids are two F1s, one *raffonei* backcross, and one hybrid which is most likely a *siculus* backcross. If we consider the sampling years, one hybrid (*siculus* backcross) was detected in 2015 out of 30 captured individuals (only 10 individuals captured in 2015 were sequenced, but none of the non-sequenced individuals showed an intermediate phenotype; this suggests a 2015 hybrid ratio = 0.033), and three hybrids (two F1s, one *raffonei* backcross) were detected in 2017 amongst a total of 88 lizards sampled from Capo Grosso (2017 hybrid ratio = 0.034).

### 3.3 Assessment of the parental species and phenotype for the genetic hybrids

We identified the maternal species of each of the four hybrid individuals by sequencing the 12S rRNA mtDNA region. Mitochondrial DNA sequences obtained for the hybrids CGI35 and CGI38 were identical to the 12S sequence of a *P. siculus* individual from Vulcano island (accession KX080573: Mendes *et al*., 2016); sequences of the hybrids CGI13 and CGI36 show a single substitution (genetic identity of 99.7%) from the sequence found in four *P. raffonei* individuals from Strombolicchio islet (KY562000: Salvi *et al*., 2017; MW619265-267: Salvi *et al*., 2021). Sequences of *P. siculus* and *P. raffonei* show 15 alternatively fixed SNPs at this 12S gene fragment (length: 348 base pairs). This indicated that the two F1 hybrid lizards originated from either a *P. raffonei* female (CGI13) or a *P. siculus* female (CGI35). The *raffonei* backcross lizard originated from a *P. raffonei* female in the backcross or in the F1-producing cross. The low-coverage sample (CGI38) originated from a *P. siculus* female.

Inspection of the phenotypes suggests hybrids more closely resembled the maternal species in both dorsal and ventral patterns, regardless of hybrid class and sex. CGI13 (F1, *P. raffonei* maternal haplotype) and CGI36 (*raffonei* backcross, *P. raffonei* maternal haplotype) had a phenotype more closely resembling the *P. raffonei* phenotype with few to many dark markings on the chin shields and throat, orange ventral colouration, a dark vertebral stripe flanked by two brown/green stripes on the dorsal surface (Figure 5A). Conversely, CGI35 (F1, *P. siculus* maternal haplotype) and CGI38 (*siculus* backcross, *P. siculus* mother) resemble a *P. siculus* phenotype with white belly and throat with pale spots and mostly reticulated dorsal pattern (Figure 5B).

**Figure 5.**
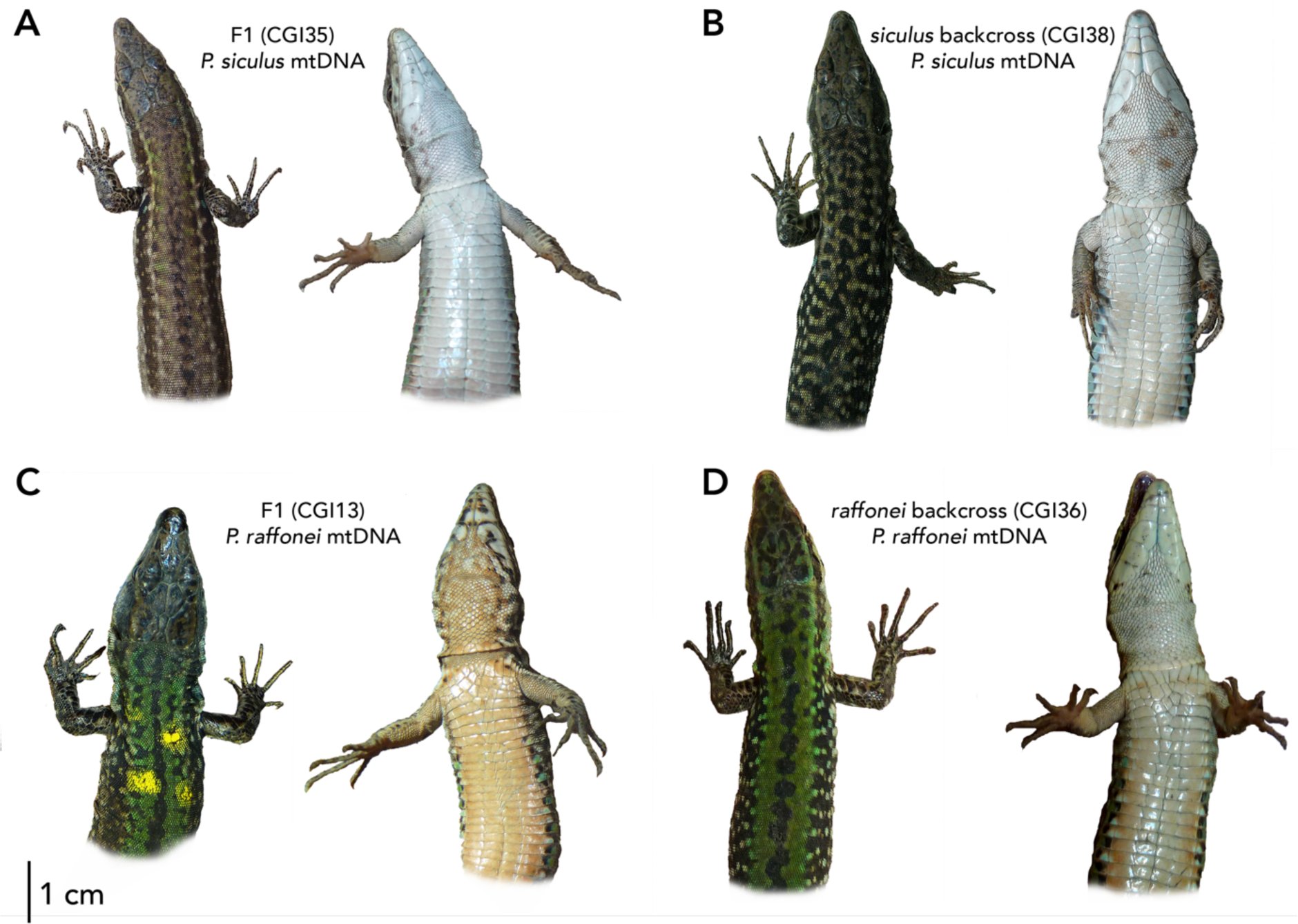
Dorsal and ventral phenotypes of the four hybrid individuals more closely resembled the maternal species, regardless of hybrid class. (A) CGI35 (F1, *P. siculus* maternal haplotype) and (B) CGI38 (*siculus* backcross, *P. siculus* maternal haplotype) resemble a *P. siculus* phenotype with white belly and throat with pale spots and mostly reticulated-blurred dorsal pattern. (C) CGI13 (F1, *P. raffonei* maternal haplotype) and (D) CGI36 (*raffonei* backcross, *P. raffonei* maternal haplotype) resemble the *P. raffonei* phenotype with orange ventral colouration, dark markings on the chin shields and throat, a dark vertebral stripe flanked by two green stripes on the dorsal surface.

### 3.4 Genetic diversity and effective population size estimates of *P. raffonei*

Estimates of genetic diversity were calculated using individuals with the lowest amount of missing data for each species (*P. raffonei*=60; N *P. siculus*=30). All the estimates of genetic diversity were lower in *P. raffonei* compared to *P. siculus* (Table 1). Estimates of the inbreeding coefficient (*F*_IS_) were slightly higher in *P. siculus*, with confidence intervals above zero. Iterative downsampling of the sample size in the *P. raffonei* dataset did not affect the results, suggesting robustness in the estimations. Estimates of the effective population size (N_E_) of *P. raffonei* from Capo Grosso were 63.8 (95% CI = 54.7 – 75.5) when including singletons, translating to an N_E_/N_C_ ratio of 0.06. Excluding singletons lead to slightly smaller N_E_ values but with larger confidence intervals (N_E_ = 49.8; 95% CI = 25.5 – 133.7).

**Table 1.** Genetic diversity estimates for the Aeolian wall lizard (*Podarcis raffonei*) from Capo Grosso and the Italian wall lizard (*Podarcis siculus*) from Vulcano. Estimates of diversity include expected heterozygosity (H_E_), observed heterozygosity (H_O_), allelic richness (A_R_), and the inbreeding coefficient (F_IS_). Values are shown for both species using the full datasets of individuals with the lowest amount of missing data (*P. siculus n* = 30 and *P. raffonei n* = 60) with numbers in brackets representing the values obtained after downsampling iterations for *P. raffonei*.

| Species | Dataset | $H_E$ | $H_O$ | $A_R$ | $F_{IS}$ |
| --- | --- | --- | --- | --- | --- |
| Aeolian wall lizard<br>(*Podarcis raffonei*) | Average and range of full dataset<br>($n=60$) and iterations ($n=30$) | 0.022<br>(0.021-0.023) | 0.022<br>(0.022 - 0.023) | 1.07<br>(1.06 - 1.08) | -0.005<br>(95% CI: -0.001 - 0.002)<br>(-0.014 - 0.001) |
| Italian wall lizard<br>(*Podarcis siculus*) | Full dataset ($n=30$) | 0.0570 | 0.0547 | 1.25 | 0.041<br>(95% CI: 0.036 - 0.046) |

## 4. Discussion

The promontory of Capo Grosso in the island of Vulcano hosts the largest remaining population of the Critically Endangered Aeolian lizard. This population represents the last stronghold for the species on a large island, where *P. raffonei* persisted after the historical introduction of *P. siculus* (Capula, 1993, 1994; Sherpa *et al*., 2023). Recent observations of lizards with an intermediate phenotype in Capo Grosso resulted in renewed concern, especially given that interspecific hybridisation with *P. siculus* could quickly drive this population to extinction (Ficetola et al., 2021). Our high-density panel of SNPs showed limited hybridisation (∼3%). Just a handful of intermediated individuals were hybrid, there is no evidence of older hybrid classes, and most individuals sampled from Capo Grosso are in fact pure *P. raffonei*. Therefore, we cautiously suggest that hybridisation does not currently represent the major threat for *P. raffonei* in Capo Grosso. These findings are in contrast with the speculation that hybridisation caused the disappearance of *P. raffonei* from Capo Grosso based on phenotypic observations (Lo Cascio & Sciberras 2020) and casts doubt on the hypothesis that hybrid swamping was the primary reason for the disappearance of the species from the Aeolian archipelago.

### 4.1 Low contemporary hybridisation between the Aeolian wall lizard and the Italian wall lizard

Our genome-wide estimate of hybridisation between *P. raffonei* and *P. siculus* suggests a low (∼3%) rate of hybridisation compared with other congeneric pairs (e.g. Caeiro-Dias *et al*., 2021). This is in contrast with a previous genetic study on Vulcano, which reported a much higher ratio of F1 hybrids (15%; Capula, 1993). This can be explained by the very small panel of loci employed by this early study (four allozymes). Assessments of hybrid zones based on a few (<100) loci can yield inaccurate estimates of introgression rate (Boecklen & Howard 1997), stressing the need for methods which extensively cover the genome. Despite the low information content in each individual SNP, the large number of markers provided by approaches such as RAD-seq offers a cost-effective method to achieve accurate hybrid classification (Twyford & Ennos, 2012) while genotyping a large number of individuals (Ravagni *et al.,* 2021). Other studies observed discrepancies in the hybridisation rate measured using different marker typologies (e.g. Dupuis & Sperling, 2016; Miralles *et al*., 2023), suggesting caution in the interpretation of hybridisation rates derived by allozyme data and in the comparison of estimates based on different marker types.

Aside from methodological differences, a further potential explanation for the discrepancy between the results here and those of Capula (1993) is that interspecific interactions on Vulcano Island have probably changed in the last ∼40 years, especially given the dramatic decline of *P. raffonei*. In 1951, *P. raffonei* was reported to be the predominant species in the northern areas of the island (Mertens, 1955). In the late 1980s, Capula witnessed the replacement of *P. raffonei* by *P. siculus*, but still found pure *P. raffonei* individuals and hybrids in these areas (Vulcanello and at the base of Gran Cratere; Capula, 1993; Capula pers. comm.), not far from Capo Grosso (∼1 km). The expansion of invasive lizards (as observed by Capula) probably led to an increased frequency of interspecific encounters and thus more frequent hybridisation. On the other hand, the distal portion of the Capo Grosso peninsula, where *P. raffonei* still occurs, is connected to the main island by a narrow and tortuous rocky isthmus (∼400 m long, minimum width ∼10 m, minimum altitude 1-2 m above-sea-level; Fig. 1) which probably represents a strong geographical filter against the invasion of *P. siculus*. The strength and type of the habitat barrier (in this case the isthmus) can have substantial effects on the flux of alleles in hybrid zones, potentially limiting introgression levels (Barton, 1979). We remain blind about the temporal movement of the hybridisation zone, but it can be hypothesised that it has moved over time and that hybridisation rate increased at the expansion front of *P. siculus*, and then decreased in extent as *P. siculus* outcompeted *P. raffonei* until near-complete replacement. The *P. siculus* lizards sampled in this study were mostly from NE Vulcano (Vulcanello), where the historical hybrids were observed, but we found no trace of introgressed alleles from *P. raffonei*. This suggests that introgression has not been the main driver of the replacement of Aeolian lizards by invasive lizards, and that other processes (e.g. competition; Capula 1992; Capula *et al*., 2002; D’Amico *et al*., 2018) may have taken place and require further investigation.

Demographic and spatial trends of these lizards have been likely determined by their different ecological response to the anthropogenic impact on native habitat. The two species show clear ecological differences, with *P. siculus* thriving in anthropic environments, including open, and urban habitats (Biaggini *et al*., 2009; D’Amico *et al*., 2018). The dispersal of *P. siculus* is limited by habitat quality and density-dependent, being higher in crowded habitats (Vignoli *et al*., 2012). Opportunistic transects along the Capo Grosso promontory indicate that *P. siculus* density is very low and decrease along the isthmus (L. Vignoli and B. Gambioli, pers. comm.), further supporting the hypothesis that the isthmus is an efficient barrier to the invasion by *P. siculus*, possibly preventing extensive hybridisation at this site. In Capo Grosso, hybridisation rate apparently remained at similar, low rates between 2015 and 2017. Still, data covering a longer timeframe are needed to ascertain the dynamics occurring in this area.

Besides the role of habitat changes and species abundance, the low rate of hybridisation could also be due to evolutionary mechanisms associated with the genetic differentiation between species and the fitness of hybrid offspring (Barton & Hewitt, 1985). The high genetic differentiation (*F*_ST_ = 0.92) supports an old divergence between *P. raffonei* and *P.* siculus (around 11-18 mya; Salvi *et al*., 2021; Yang *et al*., 2021). Nevertheless, *Podarcis* lizards are typified by pervasive ancestral hybridisation and introgression (Jančúchová-Lásková *et al*., 2015; Yang *et al*., 2021, 2022), suggesting that postzygotic reproductive isolation mechanisms are fluid. Hybrid fitness is a further point of consideration when interpreting hybridisation rates. In lizards, reduced hybrid fitness can result from changes in intraspecific competition (MacGregor *et al*., 2017), sexual selection (While *et al*., 2015), and reduced reproductive function, including the production of fewer sperm and fewer eggs (Gorman *et al*., 1971). The fitness of first-generation hybrids is presumably very low given the observation of atrophied gonads in F1 hybrids (Capula, 1993). The low fitness of later-generation hybrid offspring can be indirectly inferred from the apparent lack of later generation hybrids in our analyses. Negative effects of hybridisation are often manifested in later-generations, such as F2s and higher order hybrids (Edmands, 1999; Muraro *et al.,* 2022). Nevertheless, the detection of a few individuals with backcross genetic signatures indicate that early-generation hybrids are not always sterile as previously proposed (Capula, 1993). In a review of hybridisation between 94 pairs of genetically distinct lizard species and subspecies, the majority of F1 hybrids were also found to be fertile, thus allowing backcrosses with at least one parental species (Jančúchová-Lásková *et al*., 2015). The individual identified as a backcross with *P. raffonei* is of special concern, as even infrequent F1 hybrids can facilitate backcrossing, leading to the risk of parental genotype displacement (Rhode & Cruzan, 2005). Nevertheless, the rarity of backcrosses suggests that they have low fitness. This is further supported by the lack of introgressed alleles observed in *P. siculus* from Vulcanello (see above). Overall, the hybridisation between *P. raffonei* and *P.* siculus is a spatially complex and highly dynamic process, and strong demographic and ecological factors, including species abundance and habitat disturbance, as well as postzygotic mechanisms, probably determine hybridisation rate and extent.

### 4.2 Phenotypic identification of hybrids

The identification of which species acted as the maternal parent of hybrids may assist future studies wishing to address which genetic factors have contributed to the observed fitness outcomes (Chan *et al*., 2019). The phenotype of hybrid individuals suggests a maternal effect on the phenotype (Fig. 5; Soletchnik *et al*., 2002; Kirk *et al*., 2005), but this remains speculative given the small number of identified hybrids. In any case, the phenotypes of these hybrids are not markedly different from pure individuals, confirming the challenge of morphological identification (Capula, 1993; Ficetola *et al*. 2021). The identification of species identity and hybrids without genetic data (e.g. Lo Cascio & Sciberras, 2020) is thus unreliable, stressing the need for continuous genetic monitoring prior to any management action. Indeed, our genetic analysis corroborates that green colouration is a plastic trait of *P. raffonei*. Several *P. raffonei* individuals show a seasonal transition, shifting from the typical brown pattern into a green phenotype during spring (Ficetola *et al*., 2021). Green colouration has not been previously reported for *P. raffonei* from Vulcano by earlier studies (Capula *et al.,* 2002; Capula & Lo Cascio 2011), but seasonal dorsal colour changes have been observed in other *Podarcis* lizards including *P. siculus, P. waglerianus*, *P. carbonelli* and *P. bocagei* (Galan *et al.,* 1995; Faraone *et al*., 2006; Sá-Sousa, 2015; Pellitteri-Rosa *et al.,* 2020; Storniolo *et al.,* 2021). Further research is needed to understand the ecological and evolutionary drivers of this phenotype, and its fitness implications.

### 4.3 Current genetic status of the Aeolian wall lizard in Capo Grosso

Recent genomic analyses showed that *P. raffonei* has the lowest genetic diversity of any of the 26 *Podarcis* species, including other island-endemics (Yang *et al.,* 2021), and the lowest heterozygosity among seven species belonging to distinct squamate families (Gabrielli *et al.,* 2023). The low genetic diversity of *P. raffonei* from Capo Grosso corroborates this. Neutral genetic diversity is often used as a surrogate measure of adaptive capacity (Kardos *et al*., 2021). Nevertheless, low genetic diversity does not always prevent probable adaptation in *Podarcis* lizards (Sherpa *et al*., 2024). Furthermore, some studies suggest that populations and species can survive for long periods of time with low genetic diversity (*e.g.*, Pečnerová *et al*., 2023) if they can effectively purge genetic load (Bertorelle *et al*., 2022). Ongoing research quantifying the genetic diversity and genetic load in *P. raffonei* will further clarify how at-risk the species is regarding these harmful genetic risk factors.

The effective population size (N_E_) is one of the most important indicators of evolutionary potential (Waples 2022) and extinction risk (Antao *et al*., 2011). N_E_ of the Capo Grosso population of *P. raffonei* was small, ranging from 49.8 (95% CI 25.5 – 133.7) to 63.8 (95% CI 54.7 – 75.5), depending on the exclusion or inclusion of singleton markers, respectively.

Although comparisons to N_E_ estimates are challenged by methodology and the marker type used, the N_E_ estimate for *P. raffonei* in Capo Grosso is lower compared to other island-endemic reptiles (e.g. *Gongylomorphus bojerii*: N_E_: 99.6 – 228; (Michaelides *et al*., 2015)*; Sceloporus occidentalis becki*: N_E_: 175–236 (Trumbo *et al*., 2021), and similar to islet populations of *Podarcis gaigeae*: N_E_: 39.4 – 97.7 (Runemark *et al*., 2010). However, because N_E_ is challenged by biases, the ratio of effective size to census size, N_E_ /N_C_ is deemed a more useful indicator of the extent of genetic variation expected in a population (Frankham, 1995; Palstra & Fraser, 2012). Our estimate of N_E_ /N_C_ for *P. raffonei* is 0.05 – 0.06. This is lower than the average N_E_ /N_C_ of 0.1, estimated over a range of different animal taxa (Frankham, 1995; Palstra & Ruzzante, 2008; Palstra & Fraser, 2012), and is lower than the most recent review of N_E_ /N_C_ calculated for other reptile species (range: 0.08-73; Hoban *et al*., 2020).

The extremely low estimates of genetic diversity and effective population size probably indicate a reduced evolutionary potential of the Capo Grosso population that can determine a limited ability to withstand environmental stressors, thus increasing extinction risk (Frankham 2008). Given recent evidence of a further demographic decline in this population (L. Vignoli and B. Gambioli unpublished data), we stress the urgency of management plans, combined with robust demographic and genomic monitoring.

### 4.4 Hybridisation: good or bad for small, isolated populations?

Interspecific hybridisation is arguably one of the most controversial, and neglected, topics in conservation (Draper *et al*., 2021). Hybridisation can drive rare species to extinction through genetic swamping, via hybrid replacement of the rare form, or through demographic swamping, where the overall population growth is reduced to the production of maladaptive hybrid individuals (Todesco *et al*., 2016). Moreover, extinction risk is exacerbated if the hybrids exhibit reduced fitness relative to that of either parental species (*i.e.*, outbreeding depression (Wolf *et al.,* 2001). However, for small, isolated populations of rare species which may potentially lack the genetic variation required for adaptation (Kardos *et al*., 2021), or where variation is required to avoid the negative impacts of realised genetic load (Bertorelle *et al*., 2022), natural hybridisation may represent a source of novel genetic variation that can increase evolutionary potential (Chan *et al*., 2019; Taylor & Larson, 2019). This is supported by empirical evidence that introgressive hybridisation can be central in increasing fitness, driving rapid evolution, and improving environmental stress responses (Arnold *et al*., 2001; Grant *et al*., 2005; Richards & Hobbs, 2015; Zhang *et al*., 2020). From a conservation perspective, the role of hybridisation is controversial due to concerns about the dilution of parental species’ genetic integrity, as well as defining appropriate conservation policies for hybrid populations (Allendorf *et al*., 2001). Nonetheless, correctly defining conservation and management programs for rare species with extremely fragmented populations is arguably more pressing (Ralls *et al*., 2018). On the basis of empirical studies, the inbreeding depression threat of small populations is more urgent than the potential disadvantages of outbreeding (Edmands, 2007). Given the near impossible potential for gene flow between the four geographically isolated populations of *P. raffonei*, could low natural hybridisation with *P. siculus* represent a mechanism of population rescue for *P. raffonei*? It might be assumed that the low number of identified hybrids might exclude the possibility of genetic swamping while representing a suitable scenario for adaptive introgression of positively selected variants that could improve the genetic status of *P. raffonei*. Empirical evidence suggests that even extremely low fertility or viability of early-generation hybrids, does not prevent gene flow and the establishment of new adaptive genetic cassettes (Arnold *et al*., 1999). Our results suggest that perhaps the major concern for *P. raffonei* is not about hybridisation *per se* but rather the fact that the encounter and natural hybridisation with *P. siculus* comes at the cost of imminent competitive exclusion as observed in mainland Vulcano in the last century.

## 5. Conclusions: Implications for the conservation of the Aeolian wall lizard

Genomic data has democratised the field of population genetics and can provide crucial information for conservation and management (Hohenlohe *et al*., 2021). This first population genomic analysis of the Aeolian wall lizard provides key insights on the causes of its decline, and on possible management scenarios. The interplay between the low fitness of hybrids, alongside demographic and ecological factors explain the low rate of hybridisation, and the lack of introgressed alleles in *P. siculus* from mainland Vulcano. Together, this suggests that other processes, such as interspecific competition, probably play a stronger role in the decline of *P. raffonei*. Further studies are needed to disentangle these mechanisms, and whether they are context-dependent, *i.e.*, if *P. raffonei* can withstand the impact of *P. siculus* in specific habitat refugia. On the other hand, the very low genetic diversity and recent evidence of decline highlights the urgency of management actions to avoid the extinction of this important population. Capo Grosso is deemed the largest extant population of *P. raffonei*, still the species also survives in three tiny islets. These localities are strongly isolated, but there is very poor information on their genetic features, on their divergence, and on whether they represent Evolutionary Significant Units (ESUs). The definition of a global management plan for this Critically Endangered species thus requires integrated data, combining extensive, range-wide genetic information with detailed demographic and habitat data on each population. These analyses, together with the results presented here will provide the bases for much needed conservation projects (e.g. the EOLIZARD Life Project; https://cinea.ec.europa.eu/programmes/life_en) that are crucial for ensuring the persistence of this unique component of island biota.

## Acknowledgements

We thank Delphine Rioux (LECA, UMR UGA-USMB-CNRS, Grenoble, France), Matteo Garzia and Emanuele Berrilli for help with the DNA laboratory work and Weizhao Yang and Stephanie Sherpa for the help with initial check of RAD data. We thank Andrea Melotto, Roberto Sacchi, Stefano Scali and Leonardo Vignoli for their help during fieldwork. This study was funded by the Italian Ministry for Research (PRIN project ‘Hybrind’ – 2017KLZ3MA to DS and GFF), the Mohamed bin Zayed Species Conservation Fund (Project 162514415 to GFF) and the European Commission LIFE Programme (Project 101114121 LIFE22-NAT-IT-LIFE EOLIZARD to DS). MAC and ISR are supported by grant 28014 02/SAICT/2017 and SFRH/BD/95745/2013 from FCT, Portugal. Lizards were captured and handled under permits from the Italian Ministry of Environment (PNM-0004602, PNM-0008287, and MATTM-0037921). JRP is currently supported by funding from the European Union’s Horizon Europe research and innovation programme under the Marie Skłodowska-Curie grant agreement No. 101068395 – ‘Poly2Adapt’.

## Data Availability

RAD-seq data will shortly be available on the European Nucleotide Archive (ENA).

## Author Contributions

Daniele Salvi and Gentile Francesco Ficetola designed the study. Daniele Salvi, Gentile Francesco Ficetola, Iolanda Silva-Rocha and Miguel Angel Carretero performed the sampling. Daniele Salvi, Gentile Francesco Ficetola, and Iolanda Silva-Rocha performed the molecular biology work. Josephine R. Paris designed the analytical workflow and performed the analyses with contributions from Joan Ferrer Obiol and Daniele Salvi. Josephine R. Paris wrote the manuscript with contribution from Daniele Salvi. All authors have contributed to and approved the final version of the manuscript.

## Notes

### Competing Interest Statement

The authors have declared no competing interest.

