## Supplemental Figures for "Does hybridisation with an invasive species threaten Europe’s most endangered reptile? Genomic assessment of Aeolian lizards on Vulcano island"

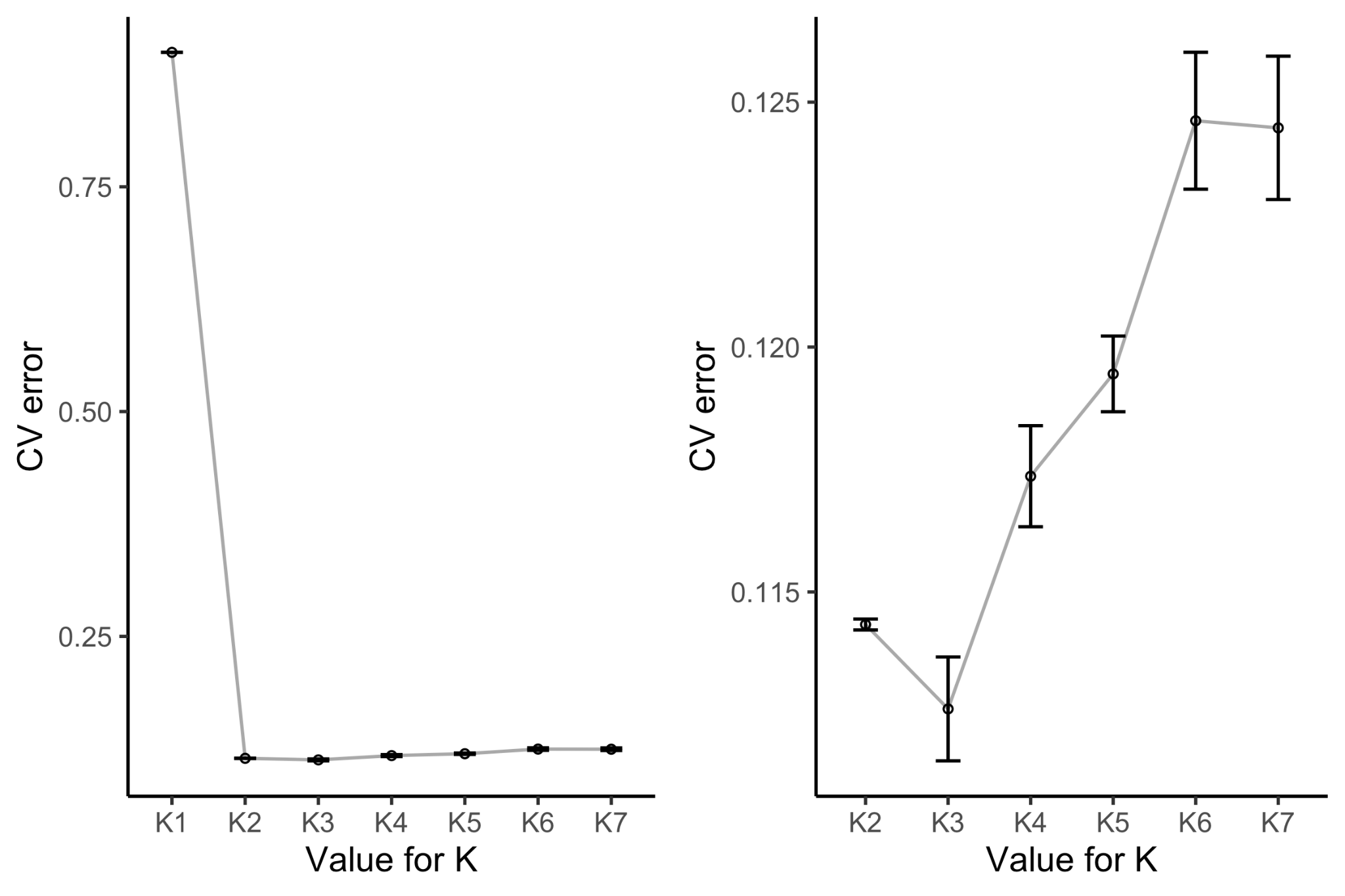

**Supplementary Figure 1**. Cross validation (CV) error rate from Admixture analysis on the full dataset comparing brown phenotype individuals from Capo Grosso (n=51), intermediate phenotype individuals from Capo Grosso (n=38), *P. siculus* individuals from Vulcano (n=35), pure *P. raffonei* individuals from Scoglio Faraglione (n=5) and pure *P. siculus* individuals from Milazzo (n=5). Both plots show the CV error for each value of *K*. Error bars represent variation across 10 independent runs. The plot on the left shows the full CV error rate, whereas the plot on the right is without *K=1*, in order to observe the higher values of *K*.

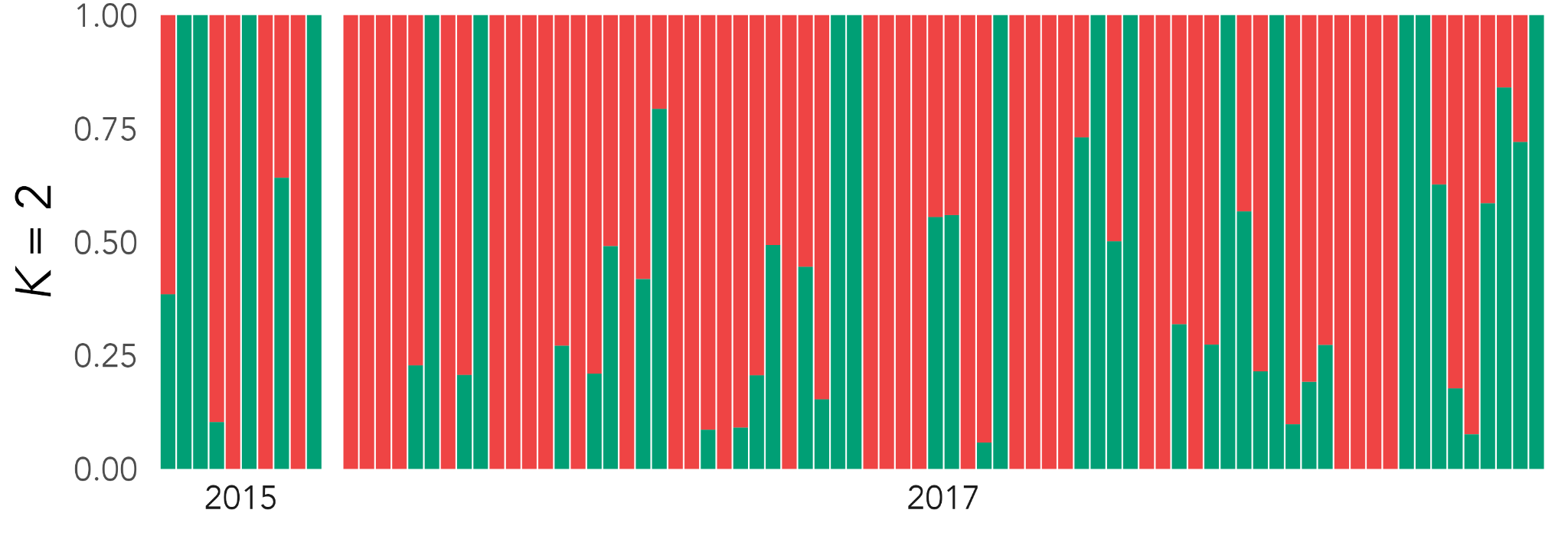

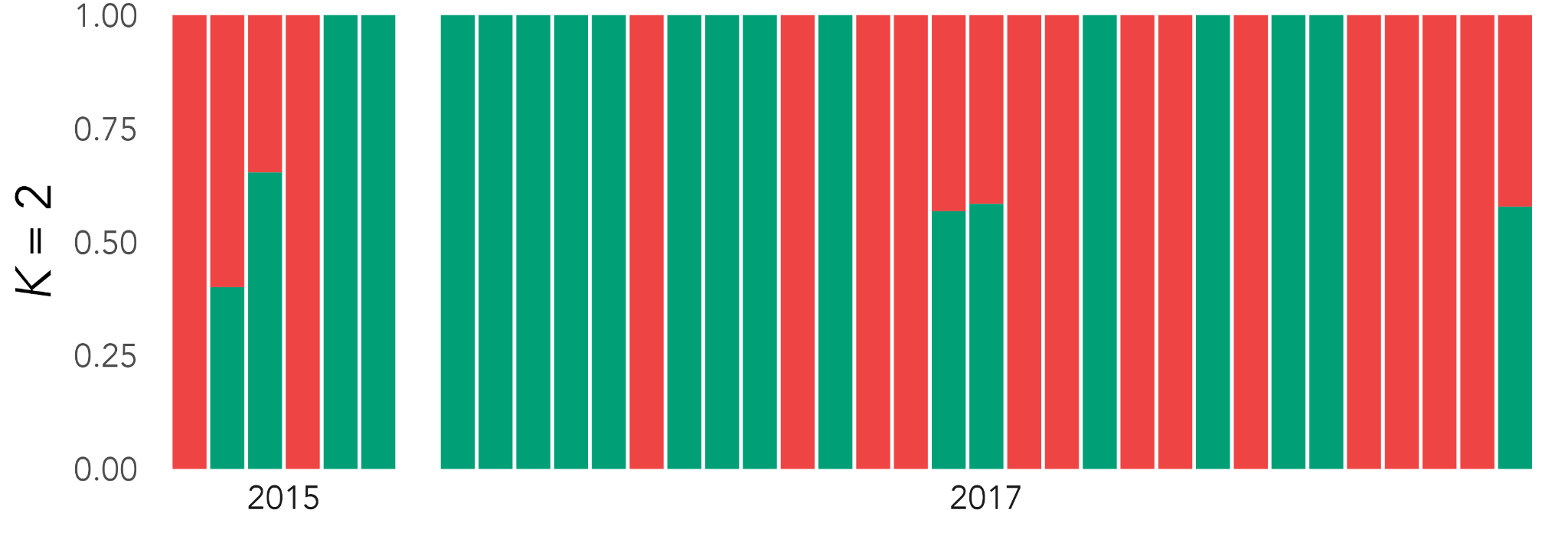

**Supplementary Figure 2.** Admixture plots showing the *K=2* proportions for *P. raffonei* (top) and *P. siculus* (bottom) to identify any potential biases arising from sampling year.

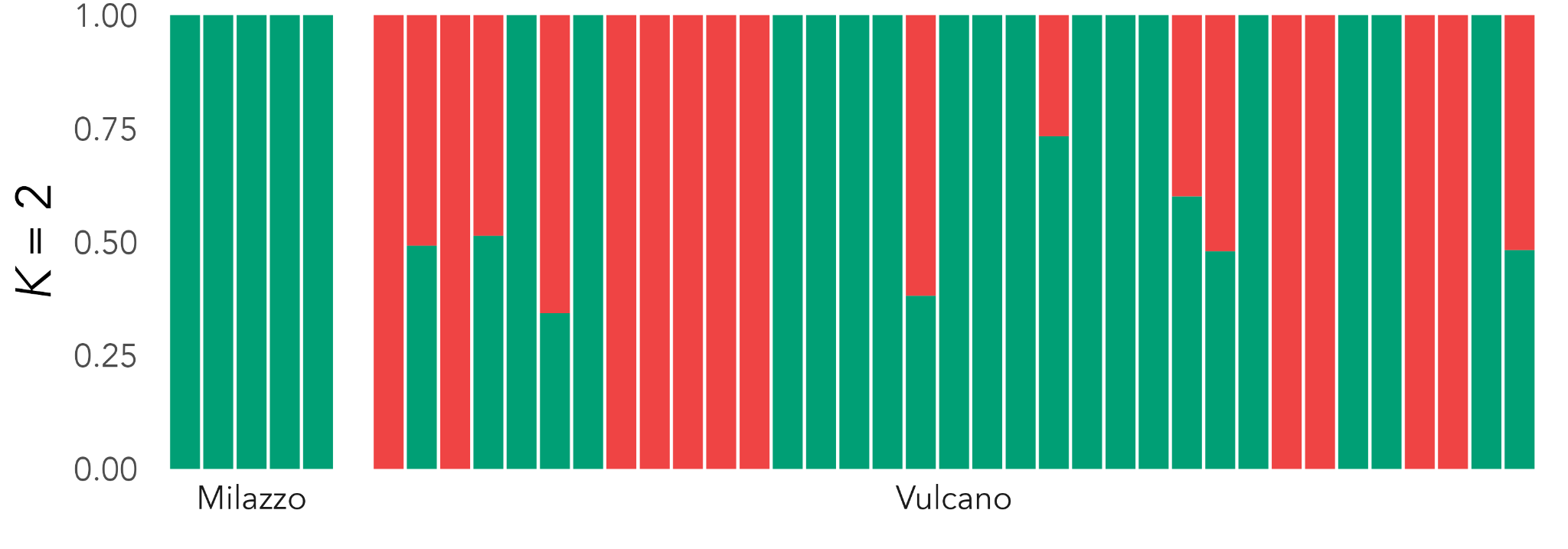

**Supplementary Figure 3.** Hierarchical Admixture analysis of *P. siculus* individuals sampled from Milazzo and Vulcano
